# Squidiff: Predicting cellular development and responses to perturbations using a diffusion model

**DOI:** 10.1101/2024.11.16.623974

**Authors:** Siyu He, Yuefei Zhu, Daniel Naveed Tavakol, Haotian Ye, Yeh-Hsing Lao, Zixian Zhu, Cong Xu, Sharadha Chauhan, Guy Garty, Raju Tomer, Gordana Vunjak-Novakovic, James Zou, Elham Azizi, Kam W. Leong

**Affiliations:** Department of Biomedical Engineering, Columbia University, NY; Irving Institute for Cancer Dynamics, Columbia University, NY; Department of Biomedical Data Science, Stanford University, CA; Department of Computer Sciences, Stanford University, CA; Department of Pharmaceutical Sciences, University at Buffalo, NY; Department of Biological Sciences, Columbia University, NY; Center for Radiological Research, Columbia University, NY; Herbert Irving Comprehensive Cancer Center, Columbia University, NY; Department of Electrical Engineering, Stanford University, CA; Department of Computer Sciences, Columbia University, NY; Data Science Institute, Columbia University, NY; Department of Systems Biology, Columbia University Irving Medical Center, NY

**Author notes:** equal contribution (Siyu He, Yuefei Zhu, Daniel Naveed Tavakol).

## Abstract

Single-cell sequencing has revolutionized our understanding of cellular heterogeneity and responses to environmental stimuli. However, mapping transcriptomic changes across diverse cell types in response to various stimuli and elucidating underlying disease mechanisms remains challenging. Studies involving physical stimuli, such as radiotherapy, or chemical stimuli, like drug testing, demand labor-intensive experimentation, hindering mechanistic insight and drug discovery. Here we present Squidiff, a diffusion model-based generative framework that predicts transcriptomic changes across diverse cell types in response to environmental changes. We demonstrate Squidiff’s robustness across cell differentiation, gene perturbation, and drug response prediction. Through continuous denoising and semantic feature integration, Squidiff learns transient cell states and predicts high-resolution transcriptomic landscapes over time and conditions. Furthermore, we applied Squidiff to model blood vessel organoid development and cellular responses to neutron irradiation and growth factors. Our results demonstrate that Squidiff enables *in silico* screening of molecular landscapes, facilitating rapid hypothesis generation and providing valuable insights for precision medicine.

## Introduction

Cells coordinate responses to environmental stimuli collectively, with variations driven by tissue heterogeneity and external cues^1,2^. Living cells function as complex, dissipative systems far from chemical equilibrium and often exhibit highly non-linear responses^3^. While single-cell sequencing enables unbiased characterization of cellular heterogeneity, predicting transcriptomic changes in responses to stimuli remains challenging. Exploring disease mechanisms and optimizing drug components require large-scale sequencing screens, which are laborious and expensive^4,5^.

Several machine-learning-based models, including scGEN^6^, scVIDR^7^, CellOT^8^, GEARS^9^, and GenePert^10^, have been developed to predict single-cell perturbation using variational autoencoders (VAE)^11^, optimal transport^8^, and graph neural networks^9^ and large language models. Despite their advances, these models face limitations in predicting high-resolution dynamic transcriptional responses, especially in transient states in organ development. Reconstructing smooth transitions of learned features remains challenging. Most models focus on specific tasks instead of a broad range of scenarios, and require both unperturbed and perturbed data as inputs, but do not fully leverage underlying biological knowledge, failing to interpolate cell states.

Diffusion models show potential in data generation through iterative refinement and learning richer data distributions than VAEs^12,13,14^. Integrating diffusion models with VAEs or performing diffusion in latent space further enhances efficiency^15–17^. In fact, these models have been applied in generating high-fidelity single-cell transcriptomic data, such as scVAEDer, scDiffusion, and scDiff^17–19^. Variants of diffusion models, such as conditional diffusional models, can manipulate smooth feature transitions and capture complex latent cellular features^19,20^. Recent perspectives highlight their potential in creating virtual cells, establishing them as a promising AI approach^21^. However, their applications in predicting gene and drug perturbations or cell development remain unexplored. Here, we introduce Squidiff (Single-cell QUantitative Inference of stimuli responses by a DIFFusion model), a computational framework designed to predict transcriptomic responses of diverse cell types to a spectrum of environmental changes, including cell differentiation, gene perturbation, and drug treatment. Squidiff is a conditional denoising diffusion implicit model^22^ generating new transcriptomes representing distinct cellular states. Squidiff also allows for the incorporation of drug compounds, including structures and dosage, when this information is available. It excels in predicting the differentiation of induced pluripotent stem cells (iPSCs) into the three germ layers, guided by stimuli vectors. Notably, Squidiff captures transient cellular states that other methods often miss. Moreover, it effectively predicts non-additive gene perturbation and cell-type-specific drug responses, as shown in glioblastoma and melanoma cells in response to novel drug combinations. The incorporation of unique drug compounds in Squidiff also enables the prediction of new drug treatment modalities.

We applied Squidiff to investigate neovascularization and vasculopathy induction in blood vessel organoids (BVOs) exposed to high-linear energy transfer (LET) radiation, uncovering potential mechanisms related to radiation-induced changes in vascular differentiation trajectories. Organoid technologies, particularly blood vessel organoids derived from human iPSCs, have shown promise in disease modeling and as drug testing platforms, especially in the context of self-organizing models that can start from a single source of stem cells^23^. This platform is useful for understanding injury and identifying potential drug candidates in extreme conditions, such as high-LET radiation exposures. In particular, these models are critical for scenarios that cannot be feasibly tested with real patients, including the risks anticipated in deep space missions. Despite the benefits of 3D systems for modeling cell types *in vitro*, the complexity of organoids and the labor-intensive nature of drug testing have limited the widespread application of these models.

To evaluate Squidiff’s capabilities in characterizing cellular dynamics in BVOs, probing the potential mechanisms of neutron radiation damage associated with secondary radiation sources in deep space, and facilitating candidate drug discovery, we applied Squidiff to predict molecular profiles of various cell compositions within BVOs throughout differentiation and development. These included endothelial cells, fibroblasts, and mural cells, the principal components of blood vessels. We validated Squidiff’s predictions with experimental single-cell sequencing data. Interestingly, Squidiff suggested a mural-to-endothelial developmental pathway consistent with recent time-series studies of vessel organoids^24^. Additionally, Squidiff predicted the effects of irradiation on various cell types by generating single-cell transcriptomic data upon irradiation, even with limited available information on irradiated cell types, and identified key affected signaling pathways at each stage of injury. Incorporating an FDA-approved radioprotective drug, G-CSF, Squidiff further predicted that G-CSF might have a protective effect by promoting vascular specification compared to irradiated groups. These findings provide a comprehensive assessment of cell heterogeneity and differentiation trajectories in vessel organoids exposed to radiation and G-CSF proteins. Squidiff thus shows the power of capturing high-resolution cell dynamics under environmental changes by leveraging the strengths of diffusion models.

## Results

### Squidiff predicts transcriptomic changes through denoising

Squidiff is a diffusion model designed to predict single-cell transcriptomic changes in response to cell state changes such as cell differentiation, development, and exposure to various physical or chemical stimuli across different cell types (**Fig. 1a**). Leveraging a neural network architecture and continuous denoising process, Squidiff predicts future and past cell states when stimuli are specified, and can further predict the effects of multiple drug combinations as well as multiple gene perturbations.

**Figure 1.**
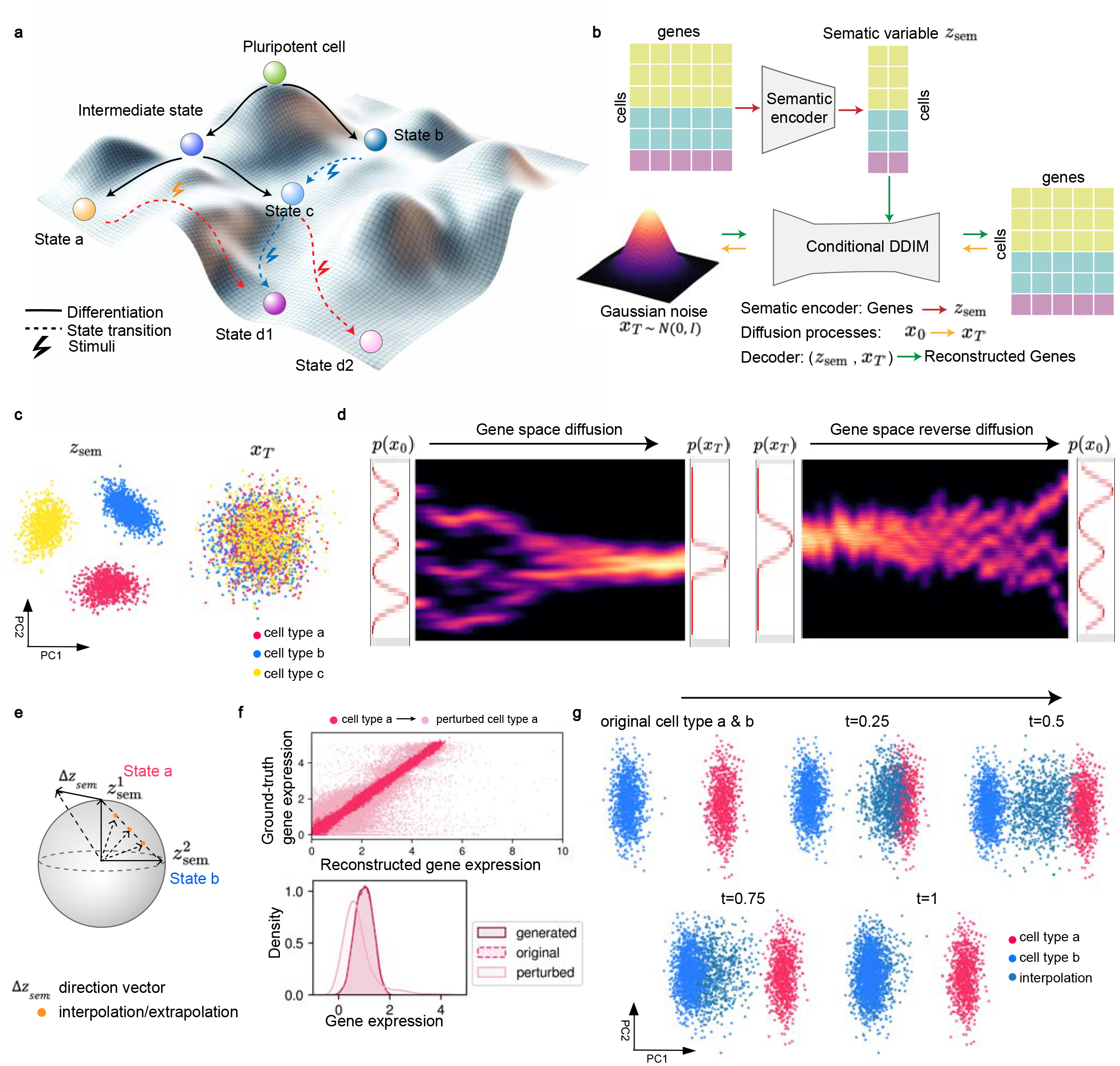
Overview of Squidiff and its performance on synthetic data. **a**, Waddington’s landscape illustration depicting cell differentiation and transdifferentiation paths from a pluripotent cell to various states (State a, b, c, d1, and d2). Solid black lines represent differentiation, while dashed lines indicate state transitions due to external stimuli. **b**, Diagram of the Squidiff model architecture, based on diffusion autoencoders^20^. The model comprises a semantic encoder and a conditional denoising diffusion implicit model (DDIM). The semantic encoder maps single-cell RNA sequencing data into a semantic latent space (*z*_*sem*_). The conditional DDIM includes a diffusion process that incrementally adds noise to input data *x*_0_, transforming it into Gaussian noise *x*_*T*_after T steps, and a denoising process that decodes the latent variables (*z*_*sem*_, *x*_*T*_) to generate gene expression profiles (See **Methods** for details). **c**, Principal component analysis (PCA) of latent representations: *z*_*sem*_ (left) shows clustering of cell types a, b, and c, while *x*_*T*_(right) displays stochastic variations across the same cell types. **d**, Visualization of the forward diffusion process (left) and reverse diffusion process (right) in gene space. Probability distributions p(*x*_0_) and p(*x*_*T*_) illustrate the transition from the original gene space to Gaussian noise and back to the gene space via denoising in the reverse process. **e**, Schematic of latent space manipulation for generating new cell states. Arrows represent direction vectors (*Δz*_*sem*_) used for interpolation and extrapolation between states, with the spherical structure indicating the semantic latent space. **f**, Top: Correlation between ground truth and reconstructed gene expression levels for cell type A and its perturbed state, showing high accuracy in predictions. Bottom: Density plot comparing the distribution of gene expression values between generated, original, and perturbed data, indicating successful reconstruction by Squidiff. **g**, PCA visualization of gene expression for cell types a and b, illustrating the temporal progression of interpolated states from *t* = 0 (original) to *t* = 1 (fully interpolated). Intermediate time points (*t* = 0.25, *t* = 0.5, *t* = 0.75) show the gradual transition between cell types a and b in the latent space.

Squidiff integrates a diffusion model for data generation and a semantic encoder for encoding latent representations^11,25^. This integration enables the generation of new transcriptomic data while modulating states via latent variables. Specifically, the semantic encoder maps single-cell transcriptomic data into a unified latent representation space comprising semantic variables (*z*_*sem*_), while the diffusion model generates target cell transcriptomes from denoising a Gaussian noise *x*_*T*_ conditioned on *z*_*sem*_ via a standard denoising process (**Fig. 1b, Supplementary Fig. 1a**, see **Methods** for details). Overall, Squidiff is capable of generating transcriptomic data reflecting cell-type variations, cell-state transitions, and cell-type-specific responses to multiple stimuli such as drug and gene perturbations.

To explain the Squidiff framework, we first show an application to synthetic single-cell RNA sequencing data generated using Splatter^26^ (**Supplementary Fig. 1b**, see **Methods**), which simulated gene expression based on a gamma-poisson distribution. We simulated single-cell transcriptomics data for three distinct cell types. During the diffusion process, iterative noise addition gradually transformed the three cell types into Gaussian noise after 1000 steps (**Supplementary Fig. 1c**). The semantic latent variable (*z*_*sem*_) captured biologically meaningful variations in gene expression associated with specific cell states or responses to stimuli, resulting in a clear separation between different conditions in the latent space (**Fig. 1c**). Meanwhile, the Gaussian noise *x*_*T*_, accounts for stochasticity in the generated single-cell data. The reverse diffusion process then denoises and reconstructs the data given the condition, successfully recovering the original transcriptomic profiles (**Fig. 1d, Supplementary Fig. 1d-e**), demonstrating the model’s ability to accurately capture and reproduce cellular states with complex gene expression distribution.

To generate new single-cell gene expression data over time and in response to stimuli, Squidiff employs two methods of latent manipulation: interpolation and addition (**Fig. 1e**). Addition involves combining the original latent representation with a perturbed direction *Δz*_*sem*_. This results in a shift in the gene expression distribution, reflecting effects of the perturbation (**Fig. 1f**).

Interpolation, on the other hand, primarily uses linear interpolation^6^, which computes intermediate points along a straight line between two vectors, offering computational efficiency. For instance, during interpolation, the blue-green cluster transitions gradually from the red to the blue cluster as time *t* increases, with linear interpolation providing a direct path (**Fig. 1g**). We further validated Squidiff’s performance across three biomedical scenarios, including cell differentiation or trans-differentiation, gene perturbations, and drug response predictions.

### Squidiff predicts cell differentiation

Squidiff can predict transcriptomic profiles of specific cell states when the transcriptome of a starting point and the latent embedding learned by Squidiff for the applied stimuli are known. This capability is particularly useful for investigating the evolution of cells or missing intermediate states in cellular differentiation and development. The approach leverages the generative capabilities of the diffusion model and its manipulation of semantic latent variables.

Applied to a public single-cell transcriptomics dataset of iPSC differentiation as an example^27^, Squidiff effectively captures the state changes from iPSCs to mesendoderm and definitive endoderm cells from day 0 to day 3 through diffusion (**Supplementary Fig. 2a**) and denoising processes. The model was trained on data only at day 0 and day 3 (**Supplementary Fig. 2b**). With the semantic encoder, the semantic latent variables *z*^0^_*sem*_ and *z*^3^_*sem*_ day 0 and day 3 represented distinct information for day 0 and day 3. The computing of *Δz*_*sem*_ by subtracting *z*^3^_*sem*_ from day 3 and *z*^0^_*sem*_ from day 0 represents the direction vector for the averaged stimuli over the period (**Fig. 2a, Supplementary Fig. 2c, Methods**). Starting with the initial states at day 0 and applying the learned stimuli direction, Squidiff accurately predicts single-cell transcriptomics from day 1 to day 3 (**Fig. 2b-c**). Additionally, applying the stimuli vector to the predicted transcriptomics on day 1 also results in an accurate prediction of cell states on day 2 (**Fig. 2c**).

**Fig. 2:**
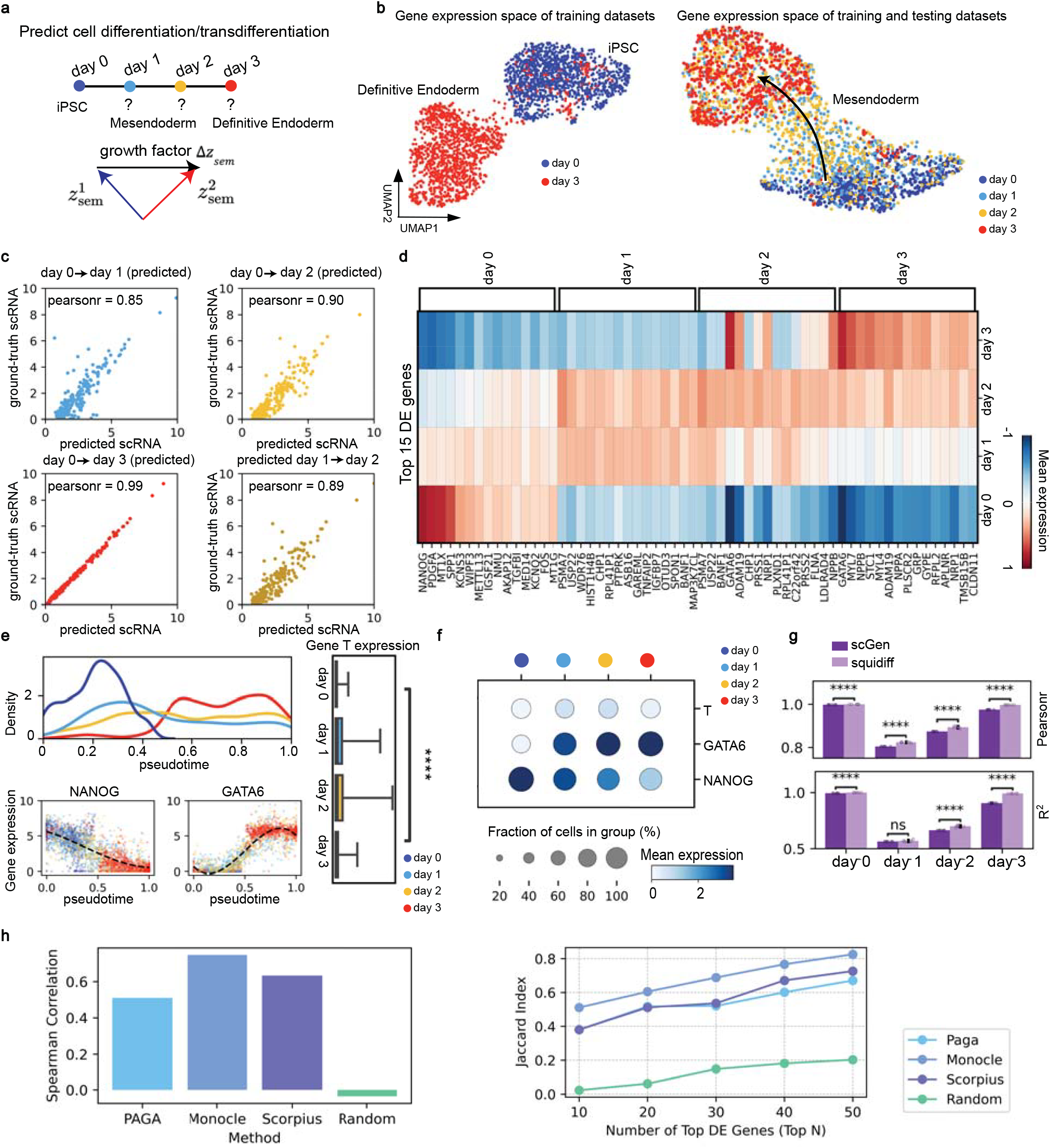
Squidiff predicts cell differentiation. **a**, Schematic of cell differentiation or transdifferentiation prediction using Squidiff. The model extracts semantic latent variables (*z*^1^_*sem*_ and *z*^2^_*sem*_) and computes the difference (*Δz*_*sem*_) to define the direction vector over the differentiation period. **b**, UMAP visualization of training datasets with iPSCs (day 0) and definitive endoderm (day 3) in blue and red, respectively. Combined training and testing datasets demonstrate differentiation from iPSCs to mesendoderm and definitive endoderm. **c**, Pearson correlation plots comparing model-predicted and ground-truth single-cell RNA-seq data across different days. High correlation coefficients indicate accurate predictions by Squidiff. **d**, Heatmap of the top 15 differentially expressed (DE) genes across days 0 to 3, illustrating the expression trends of key genes like *NANOG* and *GATA6* during differentiation. **e**, Pseudotime analysis showing cell density distributions along pseudotime (top), with changes in expression patterns of *NANOG* and *GATA6* (bottom). Comparison of inferred gene *T* expression across day 0 and 3. Box plots indicate the median (center lines), interquartile range (hinges) and 1.5× interquartile range (whiskers). n = 600 cells for each day. One-way ANOVA test was performed across days. *P* = 6.6 × 10^−56^. **f**, Dot plot showing the fraction of cells expressing *T, GATA6* and *NANOG* over different groups and days, reflecting their roles in cell state transitions. **g**, Bar plot of R-squared values and Pearson correlation comparing the prediction accuracy of Squidiff and scGen across all days. Bar plots indicate the mean (bar height) and 95% confidence interval (error bars) across six independent model runs (n=6), with individual replicate values overlaid. Two-sided independent two-sample t-test was performed. *P* = 1.97 × 10^−35^, 1.48 × 10^−9^, 5.79 × 10^−7^, 3.6 ×10^−33^ sequentially for Pearson correlation and = 3.97 × 10^−32^, 1.55 × 10^−1^, 3.93 × 10^−7^, 7.22 × 10^−35^ for R-squared values. \*\*\*\**P* < 0.0001. NS, not significant. **h**, Left: Spearman correlation between predicted cell orders from trajectory methods (PAGA, Monocle, Scorpius), random orders, and actual day-time points used in Squidiff. Right: Jaccard index of top differentially expressed genes from day 0 to day 3 across pseudotime series identified by PAGA, Monocle, Scorpius, random orders, and the actual time points used in Squidiff.

To further assess the biological relevance of the model’s predictions, we conducted differential gene expression analysis, identifying day-specific genes with distinct signatures corresponding to mesendoderm and pre-definitive endoderm stages (**Fig. 2d**). For instance, *NANOG* expression emerged as a pluripotency marker at day 0^28^, with its expression diminishing over subsequent days. In contrast, *GATA6*, which encodes a key transcription factor involved in the transition towards both mesodermal and endodermal fates^29^, showed a progressive increase. By applying pseudotime analysis (**Methods**) to the combined input data from day 0 and the model-predicted data from days 1 to 3, we observed matched fluctuations in cell proportions, with a continuous decline in *NANOG* and a corresponding rise in *GATA6* expression and signaling pathways (**Supplementary Fig. 2d**). We further examined the gene *T*, a mesodermal marker^30^, which was enriched on days 1 and 2, indicating its role in early mesoderm differentiation (**Fig. 2e-f**). These findings underscore the nonlinearity of gene expression dynamics during developmental processes and show Squidiff learns gene relationships. Notably, the latent variable embeddings from the ground-truth data of day 2 and day 3 differ from the interpolated variables, likely due to the different growth factors applied each day in this experiment. This suggests that *z*_*sem*_ represents an averaged trajectory.

Squidiff outperformed scGen^6^, a previous state-of-the-art model that combines variational autoencoders and latent space vectors for high-dimensional single-cell gene expression data (**Fig. 2g**). When trajectory inference methods such as PAGA^31^, Scorpius^32^, and Monocle^33^ were applied to the Squidiff output transcriptomics, we observed successful reconstruction of the dataset along a continuous pseudotime that closely matched the real discrete timepoints (days), surpassing random baselines. Moreover, there was a significant overlap between the genes identified based on real-time conditions and those predicted by these trajectory models (**Fig. 2h**).

### Squidiff predicts gene and drug perturbation

Squidiff also predicts cellular responses to gene perturbations and drug treatments. By leveraging vector operations, a trained Squidiff model generalizes single-cell transcriptomic data across conditions, accurately modeling complex cellular responses.

For non-additive gene perturbations, where genes interact synergistically to produce effects beyond simple additive approaches, Squidiff assumes that perturbations involving two genes can be represented as the sum of two learned semantic variables (**Fig. 3a**), allowing prediction of transcriptomic changes in wild-type cells. Specifically, we tested Squidiff on K562 cells perturbed by *ZBTB25* and *PTPN12*^*34*^, a known non-additive case discussed by GEARS^9^ (**Supplementary Fig. 3a**). Unlike GEARS, which uses graph-based prior knowledge, Squidiff requires no explicit graph structure, yet achieves highly accurate predictions (**Fig. 3b, Supplementary Fig. 3b-d, Methods**). Compared to GEARS and scGen, Squidiff consistently yields more robust and precise predictions (**Fig. 3c, Supplementary Fig. 3e**), showing its ability to capture gene perturbation effects and underlying molecular mechanisms.

**Fig. 3:**
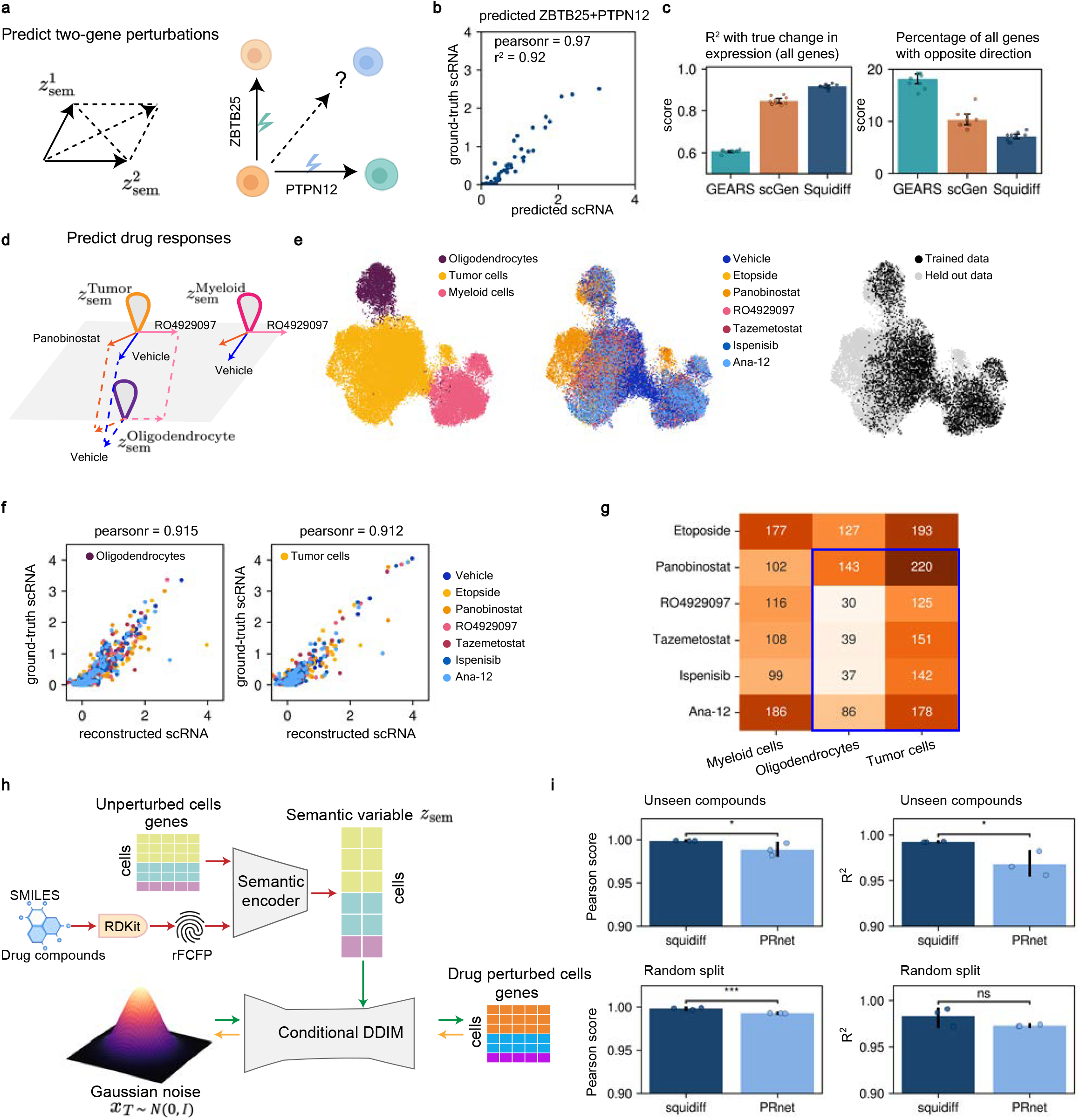
Squidiff predicts gene and drug perturbation. **a**, Schematic representation of Squidiff’s approach for predicting the effects of two-gene perturbations by leveraging semantic latent variables (*z*^1^_*sem*_ and *z*^2^_*sem*_). This method enables the exploration of combined effects of gene perturbations, such as the combination of *ZBTB25* and *PTPN12*. The cell icons were created with BioRender.com. **b**, Comparison of mean gene expression between Squidiff’s predictions and real perturbed data, evaluated using *R*^2^ values and Pearson correlation scores. **c**, Left: bar plots comparing the true expression changes induced by *ZBTB25*+*PTPN12* perturbation on control single-cell data across all genes by Squidiff, scGen, and GEARS in terms of *R*^2^ values. Right: fraction of genes where predicted expression changes are opposite to the ground truth. Bar plots indicate the mean (bar height) and 95% confidence interval (error bars), with individual data points (n=10) overlaid for each bar. **d**, Schematic representation of Squidiff’s approach to drug response prediction, where semantic latent variables (*z*_*sem*_) are manipulated based on cell types and drug treatments to model transcriptomic changes. **e**, UMAP visualization of predicted drug responses, colored by three cell types, drug treatment, and training vs. test dataset. **f**, Mean gene expression between Squidiff predicted and experimentally perturbed data, assessed using Pearson correlation scores. **g**, Heatmap displaying the response of different cell types (myeloid cells, oligodendrocytes, and tumor cells) to various drug treatments. The intensity values indicate the number of differentially expressed genes identified in each condition. **h**, Schematic diagram of Squidiff integrated with drug-rescaled Functional Class Fingerprints (*rFCFP*). **i**, Performance (Pearson correlation and R-squared scores) of Squidiff and PRNet in out-of-distribution perturbation scenarios, including both unseen compound split and random split in the sci-plex datasets. Bar plots indicate the mean (bar height) and 95% confidence interval (error bars), with individual data points (n=3) overlaid for each bar. Results are shown across three independent data splits per condition, with statistical significance assessed using a one-sided t-test. *P* = 3.58 × 10^−2^, 1.67 × 10^−2^, 9.96 × 10^−4^, = 7.46 × 10^−2^, sequentially. \*\*\*\**P* < 0.0001. NS, not significant.

Besides gene perturbations, we evaluated Squidiff on drug screening. Unlike gene targets, drug effects are often complex, involving influences on multiple genes concurrently and the regulation of associated pathways. Thus, a model predicting the overall cellular response to drugs is valuable for exploring drug mechanisms and identifying the most effective drugs with minimal side effects. We first tested Squidiff’s ability to predict the effects of two-drug combinations on cell transitions. The assumption is that the latent space captures additive or synergistic interactions implicitly. Due to the limited availability of single-cell RNA sequencing data related to drug combinations, we evaluated Squidiff on an alternative dataset 4i^8^, which profiles the molecular and morphological properties of melanoma cells treated with various drug compounds using a novel microscopic technology. We withheld drug combination data and trained Squidiff using only control cells and cells treated with single drugs, including trametinib, panobinostat, dabrafenib, erlotinib, and midostaurin. We then used Squidiff to generate single-cell data for drug combination treatments and compared the Pearson correlation between predicted and ground-truth responses from the 4i dataset. While 4i does not contain single-cell transcriptomic data and provides only ∼50 features per condition, it serves as a validation of Squidiff’s ability to generalize to drug perturbations (**Supplementary Fig. 3f**). However, the limited cellular features in this dataset may impact Squidiff’s ability to fully capture complex drug interactions compared to transcriptomic data.

We next applied Squidiff to a study investigating drug responses in glioblastoma, focusing on distinct effects of etoposide and panobinostat on tumor cells (**Fig. 3d, Supplementary Fig. 4a**). We trained the Squidiff model using a dataset consisting of randomly selected myeloid cells treated with six different drugs, as well as tumor cells and oligodendrocytes that were only exposed to etoposide (**Fig. 3e, Supplementary Fig. 4b**). Despite this constraint, Squidiff accurately predicted the effects of all six drugs on tumor cells and oligodendrocytes, demonstrating its ability to infer transcriptomic changes induced by drug treatments (**Fig. 3f**). Notably, Squidiff also identified panobinostat as having the most potent effect on tumor cells compared to other cell types among all screened drugs, consistent with the finding from the reference (**Fig. 3g**)^35^. Validating these predictions, we visualized the UMAP embedding of cells with and without treatment and observed the distinct effects of etoposide and panobinostat as previously reported (**Supplementary Fig. 4c**).

When Squidiff encounters a completely unseen drug, prediction is limited. Unlike cell type transfer learning, where Squidiff can generalize across different cell types, predicting the response to an entirely new drug is challenging, as the model has never been exposed to its molecular characteristics during training. Therefore, we integrated a drug compound adapter inspired by PRNet^36^ (**Fig. 3h**). Specifically, we incorporated the drug-rescaled Functional Class Fingerprints (*rFCFP*) from PRNet, which updates the Squidiff semantic latent variable from *z*_*sem*_ to *z*_*sem*_′ = *Enc*(*x*_0_, *rFCFP*) (**Methods**). This adaptation allows Squidiff to incorporate unseen drug information by leveraging its structural and functional properties, thereby enhancing its predictive capabilities beyond drugs included in the training set. We benchmarked Squidiff against PRNet on the sci-plex3 datasets^37^ using both an unseen drug split and a random data split. Our results show that Squidiff performs comparably to or better than PRnet (**Fig. 3i**).

### Squidiff predicts blood vessel organoids differentiation

To evaluate Squidiff’s prediction of the cascade of transcriptional programs during continuous cell state transitions and their alteration with stimuli, we employed organoid technology. Organoids, differentiated from human induced pluripotent stem cells, have emerged as unique 3D biomimetic systems for modeling human tissue development^38^, disease, and drug responses, including those for brain, liver, and lung organoids, among others^39^. However, as self-organized systems, organoids are inherently heterogeneous, and the interactions among cells within them are complex^39^. Single-cell and spatial RNA sequencing have profiled these heterogeneities in differentiation trajectories, but fully characterizing developmental processes at any given time point remains challenging. While pseudotime analysis of single-cell RNA sequencing provides snapshots of organoid development at specific stages, real-time profiling across organoids is challenging due to labor-intensive workflows and technical expertise requirements for manipulation of these engineered models^39,40^, in addition to the costly sequencing steps. Squidiff addresses these limitations by predicting cell transcriptomic changes during development and in response to stimuli.

Using BVOs as a case study for applying Squidiff, organoids were differentiated from human iPSCs and guided toward endothelial and vascular lineages, forming vascular network-like structures as early as Day 6 (**Supplementary Fig. 5a)**. BVOs show promise in modeling human vascular development and disease and serve as a powerful platform for drug discovery of human blood vessels^24^.

The fate and state transition of cells in blood vessel organoids have been studied in real-time^24^, providing an ideal dataset for comparison. In these studies, progenitor states in early BVOs were observed to bifurcate into endothelial and mural fates, highlighting the potential of BVO mural cells to differentiate into endothelial lineages (**Fig. 4a**). To test if Squidiff could replicate these findings, we generated BVOs from healthy human iPSCs to create a training dataset. We fixed a subset of organoids, and with another dissociated the organoids into single cells, performing single-cell RNA sequencing on day 11. The BVOs were exposed to differentiation processes from day -1, and progressed to mesoderm aggregates from day 3 to day 5, with vascular formation starting on day 6. We identified fibroblasts, mural cells, endothelial cells, and progenitor cells in the organoids on day 11 (**Fig. 4b, Supplementary Fig. 5a**).

**Fig. 4:**
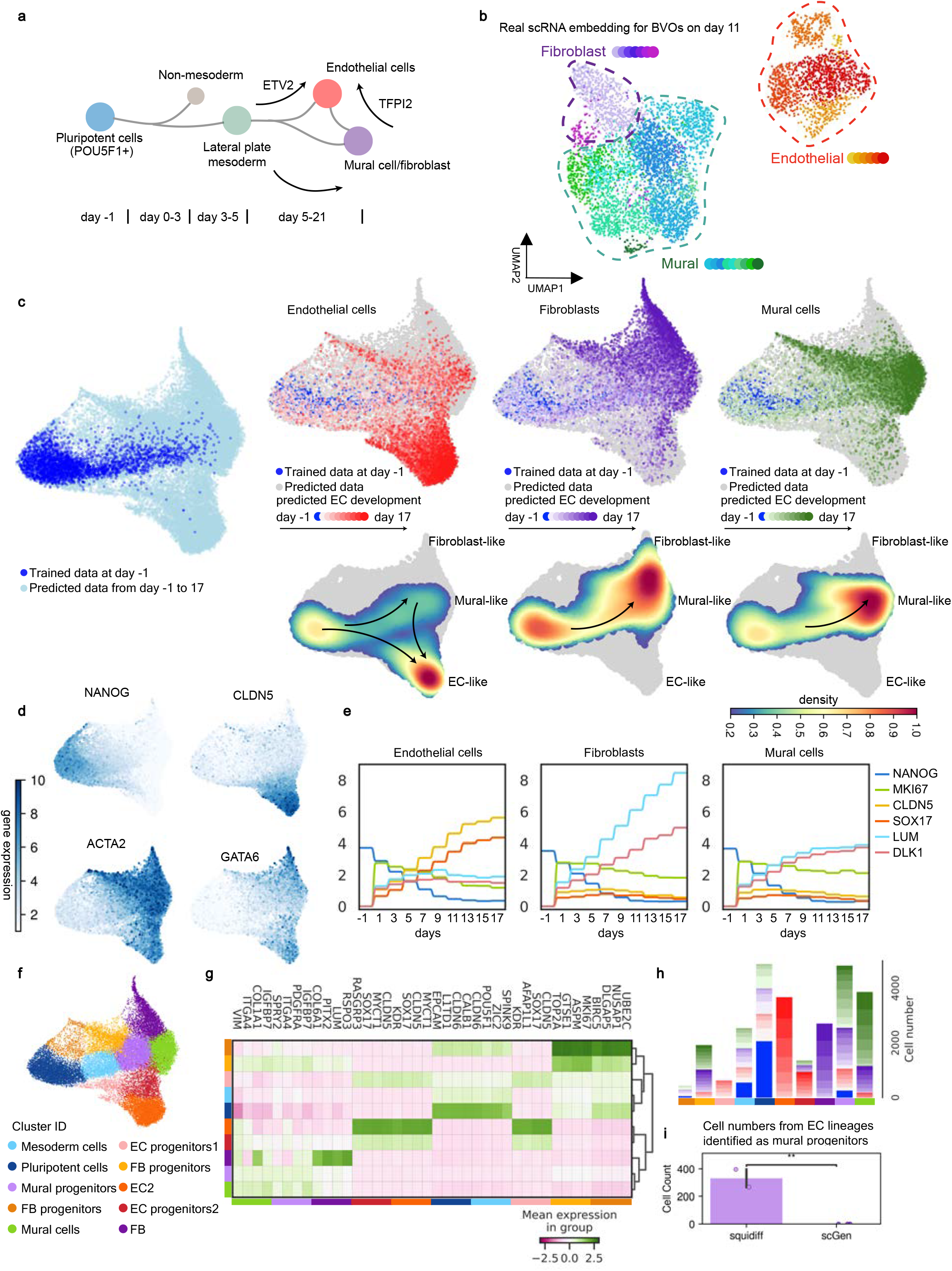
Squidiff predicts cell differentiation processes in blood vessel organoids. **a**, Diagram for hypothesized differentiation processes of iPSC into cell components in BVOs. **b**, Embedding of single-cell RNA sequencing of blood vessel organoids on day 11, colored by subtypes of endothelial cells, fibroblasts, and mural cells. **c**, Joint UMAPs of trained scRNA-seq data on day -1, namely the iPSC cells and the predicted scRNA-seq data from day -1 to day 17, with two days intervals with the semantic meaning, which was learned by only endothelial cells at day -1 and day 11. **d**, Joint UMAPs of trained scRNA-seq data on day -1 and the predicted scRNA-seq data from day -1 to day 17, colored by the markers including pluripotent genes (*NANOG, MKI67*), endothelial genes (*CLDN5*, and *SOX17*), fibroblast and mural specific genes (*COL1A1, LUM*, and *ACTA2*), and the markers denoting differentiating into mesendoderm layer (*GATA6*). **e**, Stepwise expression of markers along with days in fibroblasts, endothelial cells, and mural cells. **f**, Joint UMAPs of trained scRNA-seq data on day -1 and the predicted scRNA-seq data from day -1 to day 17, colored by cell types. **g**, Heatmap of differential gene analysis for clustering of joint trained scRNA-seq data and predicted data. **h**, Stacked barplot of the number of cells colored by the major types and days across cell type clusters. **i**, Number of endothelial cell (EC) lineages in mural progenitor and mural cell clusters, comparing Squidiff and scGen. Bar plots indicate the mean (bar height) and 95% confidence interval (error bars), with individual data points (n=2 and 3 for Squidiff and scGen) overlaid for each bar. A two-sided independent two-sample t-test was performed. *P* = 1.02 × 10^−3^.

To predict cell fates and states during blood vessel development, we combined single-cell RNA data from iPSCs and BVOs at day 11 and trained the Squidiff model on these datasets (**Supplementary Fig. 5b-c**). We then performed interpolation for days 1, 3, 5, 7, 9, 13, 15 and 17 using Squidiff. The predictions captured the differentiation and development trajectories of endothelial and mural cells from iPSCs, as shown in the distribution of marker gene expressions (**Fig. 4d-d, Supplementary Fig. 6a**). The predicted data revealed intermediate states where some endothelial cells exhibited properties akin to mural cells, while mural and fibroblast cells appeared to derive from mural progenitor cells, consistent with published findings that experimentally sequenced real-time BVOs (**Fig. 4c**). To quantify this, we tracked the expression of marker genes across days for endothelial, fibroblasts, and mural cells, identifying fluctuations in mural-specific genes, such as *LUM* and *DLK1* (**Fig. 4d**). These genes showed an increase before day 7 and a decrease after day 9, while they continued to rise in fibroblast and mural cells (**Fig. 4e**). A heatmap of differentially expressed genes across cell type clusters on the UMAP further confirmed cell type distinctions (**Fig. 4f**), and indicated the cell state transitions from iPSC to mature cell lineages (**Fig. 4g**). We observed a substantial proportion of endothelial cells in the mural progenitor population, which resembled the results of real single-cell RNA data^24^ (**Fig. 4h**). The existence of differentiating EC cells from day 5 to day 9, shown as mural progenitors, suggests the potential of ECs differentiated from mural-like cells, a feature that scGEN methods failed to capture (**Fig. 4i, Supplementary Fig. 6b-g**). This underscores Squidiff’s enhanced capability in predicting cell heterogeneity and transient cell states, offering insights for in silico modeling.

### Neutron irradiation disrupts structure and metabolism

After demonstrating Squidiff’s ability to predict gene and drug perturbations, we next evaluated its capacity to predict cellular response to physical perturbations, such as irradiation. Ionizing radiation, from radiotherapy, nuclear accidents, or deep space travel, poses serious health risks^41^, impacting not only cancerous but also non-cancerous cells, particularly endothelial, blood and immune, and local parenchymal cells within the radiated area^42^. High-energy particles like neutrons penetrate tissues and cause cellular and molecular damage^43^. Understanding radiation effects helps to develop effective countermeasures and treatments for radiation-induced damage^44^. Degenerative vascular changes are primary radiation health risks to astronaut crews on exploration missions^45^. Microvascular endothelial cells are susceptible to ionizing radiation, and radiation-induced alterations in endothelial cell function are critical factors in organ damage through endothelial cell activation, enhanced leukocyte-endothelial cell interactions, increased barrier permeability, and initiation of apoptotic pathways^42,46^. BVOs provide an excellent model for studying these effects.

On day 5 of BVO development, we exposed the cultures to either neutron or photon radiation and continued BVO culture until day 11 (**Fig. 5a**). On day 11, single-cell RNA sequencing profiled both healthy and irradiated BVOs. We identified endothelial cells, mural cells, and fibroblasts. Reduced overlap between irradiated and control cells indicated substantial molecular alterations (**Fig. 5b**). Squidiff encoded radiation exposure as a vector in the semantic space, enabling the generation of perturbed transcriptomes for different cell states in response to radiation.

**Fig. 5:**
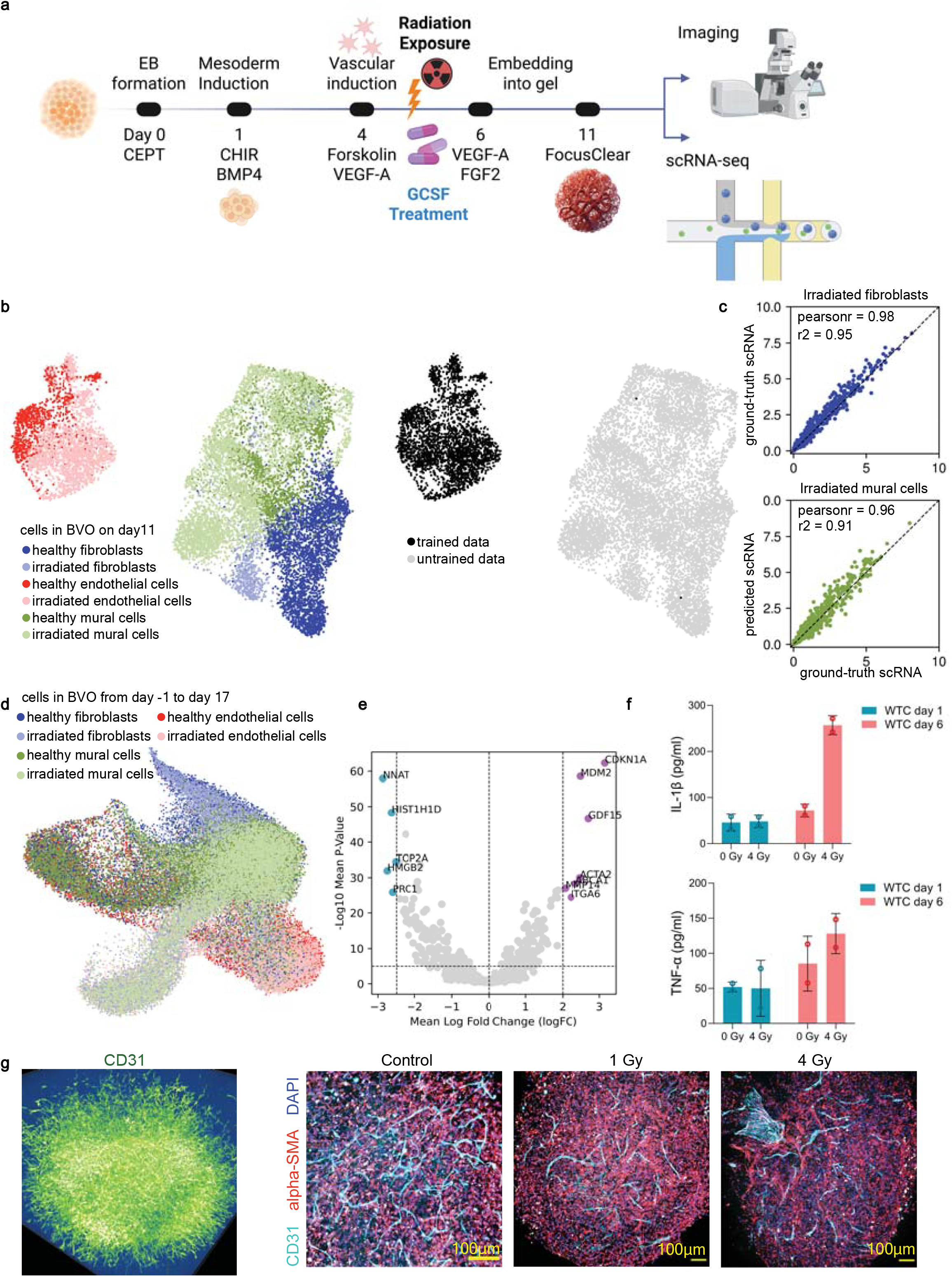
Structure damage and metabolic phenotypes alteration in blood vessel organoids induced by neutron irradiation. **a**, Experimental workflow for differentiating BVOs from human pluripotent stem cells. Key steps include embryoid body (EB) formation, mesoderm induction, vascular induction, embedding into the gel, and subsequent neutron irradiation or G-CSF treatment. Imaging and single-cell RNA sequencing (scRNA-seq) were performed on day 11. The workflow was created with BioRender.com. **b**, UMAP visualization of single-cell RNA-seq data from BVOs on day 11, showing different cell types: healthy fibroblasts, irradiated fibroblasts, healthy endothelial cells, irradiated endothelial cells, healthy mural cells, and irradiated mural cells. Trained and untrained data sets are indicated. **c**, Scatter plots showing the correlation between ground-truth scRNA-seq data and Squidiff-predicted data for irradiated fibroblasts (top) and irradiated mural cells (bottom), with Pearson correlation coefficients and R-squared values. **d**, UMAP visualization of single-cell RNA-seq data from BVOs across different developmental days (day -1 to day 17), showing the distribution of healthy and irradiated cell types over time. **e**, Volcano plot of differential gene expression in BVOs after 4 Gy neutron irradiation (30,000 irradiated cells and 30,000 control cells across day -1 to 17). Two-sided Wilcoxon rank-sum tests were performed. P values were adjusted by the Benjamini-Hochberg method. Genes exceeding logFC > 2 or < –2.5 and adjusted p value < 1e–5 are highlighted and labeled. **f**, ELISA results showing the levels of IL-1β and TNF-α in the culture medium of BVOs at day 1 and day 6, under different irradiation conditions (0 Gy, 4 Gy). Bar plots indicate the mean (bar height) and standard error of the mean (error bars), with individual data points overlaid for each bar (n=2 biologically independent replicates per group). **g**, Left: 3D rendering of a CD31-stained BVO. Right: Immunofluorescence images of BVOs showing structural changes under control conditions, 1 Gy, and 4 Gy photon irradiation. CD31 (endothelial cell marker) in cyan, alpha-SMA (smooth muscle cell marker) in red, and DAPI (nuclear stain) in blue. Scale bar, 100 *μ*m.

For validation, we masked mural cells and fibroblasts entirely during Squidiff training, using only endothelial cells under irradiation and control conditions (**Fig. 5b**). The model successfully generated transcriptomic data under irradiation (**Fig. 5c, Supplementary Fig. 7a-b**). The model was then applied to interpolate irradiated transcriptomes for endothelial, mural, and fibroblasts from day -1 to day 17. UMAP indicated that early-stage cells were most affected (**Fig. 5d, Supplementary Fig. 7c-d**), likely due to immature tissue structure and weak cell interactions. The ranking of the most differentiated genes upon irradiation across cells varied across developmental stages (**Supplementary Fig. 7e-f**).

Intriguingly, genes behaved differently over days and across cell types. For instance, *MYCT1* in endothelial cells decreased in the middle period and resumed at the latest stage of development, while *MYCT1* remained one of the most differentiated genes in mural cells and fibroblasts (**Supplementary Fig. 7e**). The most affected genes upon irradiation across all these cells included upregulation of genes such as *CDKN1A, MDM2, GDF15, ACTA2, ABCA1, MMP14*, and *ITGA6*, and downregulation of genes such as *NNAT, HIST1H1D, TOP2A, HMGB2*, and *PRC1* (**Fig. 5e**). These genes are involved in cellular responses to ionizing radiation, cell motility, DNA damage through P53 pathways, apoptosis, TNF-α signaling via NF-*κ*B, KRAS pathways, protein kinase B signaling, and inflammation responses (**Supplementary Fig. 8a-b**).

Notably, *CDKN1A* encodes p21/WAF1/CIP1, known for its role in cell cycle regulation. *MDM2* encodes a protein functioning as a negative regulator of the p53 tumor suppressor, thus playing a critical role in controlling cell proliferation and apoptosis^47^. Live-dead assays showed that the percentage of dead cells was much higher in irradiated BVOs (**Supplementary Fig. 9a-c**), consistent with radiation-induced apoptosis. BVOs decreased in size after neutron irradiation. We found that the culture supernatant post-irradiated was enriched in IL1-β and TNF-alpha, indicating enhanced induction of inflammation (**Fig. 5f**), consistent with the increased expression of *GDF15*, a stress response gene associated with the regulation of inflammation and cell survival (**Supplementary Fig. 9d**). These metabolic changes overall resulted in downstream failures in vascular formation and abnormal distribution patterns in BVOs (**Fig. 5g**), causing earlier vascular sprouting in the radiation BVOs versus controls. Overall, Squidiff effectively generated data with biological relevance and predicted the complex molecular responses to radiation exposure.

### G-CSF mitigates radiation damage in blood vessel organoids

Given disruption of vascular structures, apoptosis of cells triggered by the p53 pathways, and increased protein kinase B signaling, we tested whether the FDA-approved radioprotective drug, granulocyte colony-stimulating factor (G-CSF), could rescue defects in vasculogenesis *in vitro*. G-CSF is a glycoprotein that plays a crucial role in the production, differentiation, and function of granulocytes, a type of white blood cell essential for fighting infections^48^. It has been used in clinical settings to stimulate the bone marrow to produce more white blood cells, particularly after chemotherapy or bone marrow transplants. Recent studies suggest that G-CSF may also have protective effects on vascular tissues, promoting endothelial cell survival and enhancing tissue repair processes^49^. However, the study of the G-CSF effect on fibroblasts and mural cells in the vasculature remains limited. A known effect of G-CSF is its activation of MEK1/2, PI3K/Akt and NF-*κ*B pathways (**Fig. 6a**)^50–52^, consistent with findings on the regulation of NF-*κ*B and kinase B signaling on radiation injury (**Supplementary Fig. 8a-b**). Squidiff offers a way to predict G-CSF’s effect on other cell types when only one cell type (e.g., endothelial) responses are known. We sequenced the single-cell RNA expression in G-CSF-treated BVOs and considered it as ground truth data.

**Fig. 6:**
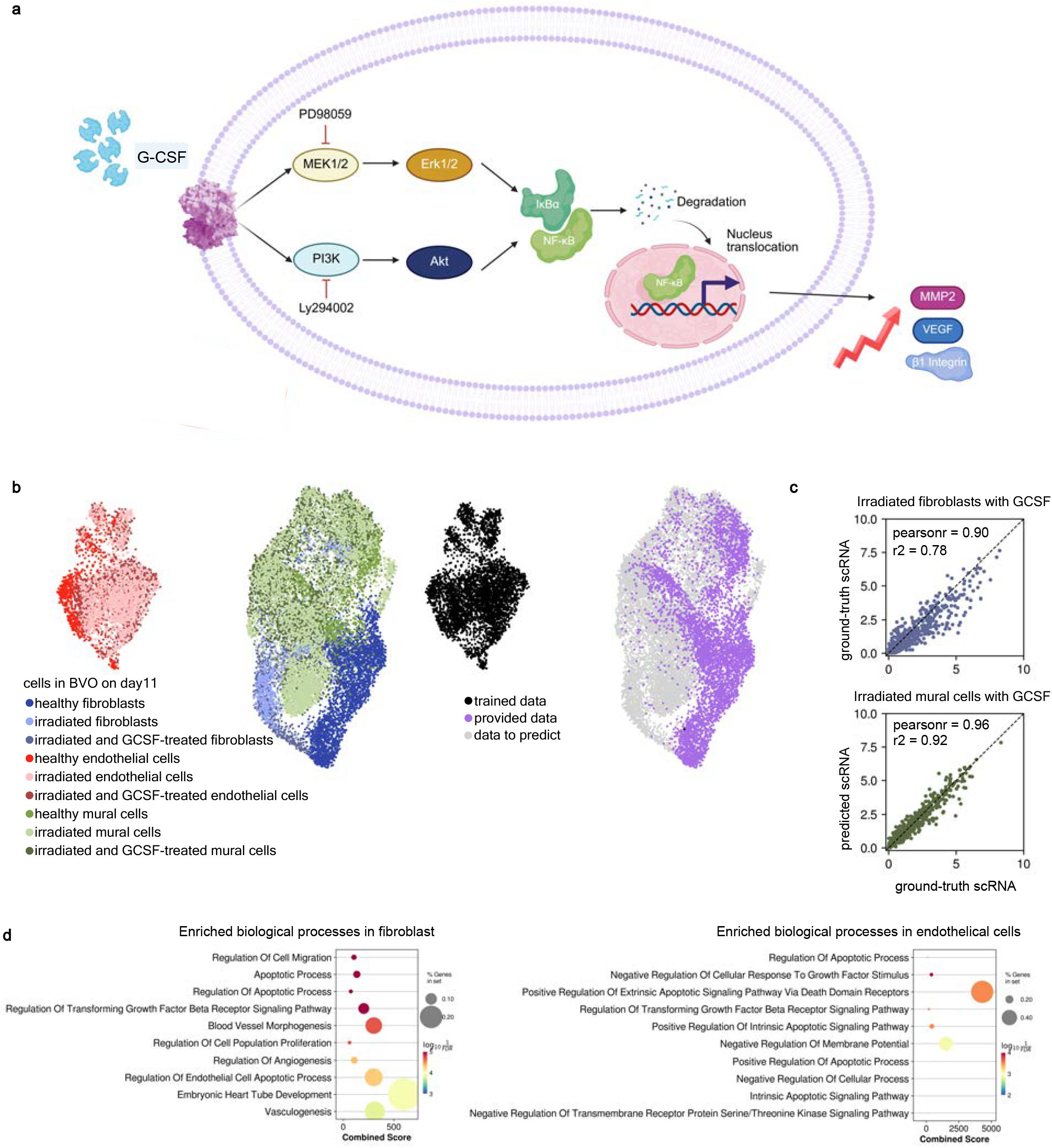
Treatment potentials of G-CSF in securing against radiation disruption in blood vessel organoids. **a**, Schematic illustration of the signaling pathways activated by G-CSF treatment^50^. G-CSF binds to its receptor (G-CSF-R), triggering downstream signaling cascades involving MEK1/2, Erk1/2, PI3K, and Akt. This leads to the degradation of I*κ*B*α* and nuclear translocation of NF-*κ*B, promoting the expression of target genes such as *MMP2, VEGF*, and *β*1 integrin, which are involved in cell survival, proliferation, and migration. The illustration was created with BioRender.com. **b**, UMAP visualization of single-cell RNA-seq data from BVOs on day 11, showing the distribution of healthy, irradiated, and G-CSF-treated cell types. The trained data includes healthy fibroblasts, endothelial cells, and mural cells. Provided data includes irradiated fibroblasts, endothelial cells, and mural cells. Data to predict includes irradiated and G-CSF-treated fibroblasts, endothelial cells, and mural cells. **c**, Scatter plots comparing ground-truth scRNA-seq data with Squidiff predicted data for irradiated fibroblasts (top) and irradiated mural cells (bottom) treated with G-CSF. The Pearson correlation and R-squared values indicate high prediction accuracy. **d**. Enriched biological processes found in fibroblasts and endothelial cells.

Given the training data on endothelial cells only (**Fig. 6b**), Squidiff successfully generated predicted single-cell transcriptomic data for fibroblasts and mural cells (**Fig. 6c, Supplementary Fig. 10a**). To validate predictions, we conducted differential gene expression analysis comparing real irradiated data with predicted G-CSF-treated data. Notably, we identified distinct sets of differentially expressed (DE) genes for each cell type, with some genes shared across multiple types (**Supplementary Fig. 10b-c**). The biological processes identified for enriched genes revealed unique roles for each cell type in response to G-CSF treatment. Fibroblasts were associated with blood vessel morphogenesis and vasculogenesis, indicating their critical role in structural development (**Fig. 6d, Supplementary Fig. 10d**). Endothelial cells showed enrichment in pathways regulating apoptosis and the cell cycle, highlighting their involvement in maintaining cell survival and proliferation (**Fig. 6d**). Mural cells, in contrast, were linked to pathways related to genomic stability, cell cycle progression, and mitosis, indicating their role in maintaining genomic integrity. This aligned with reduced cell death percentages in G-CSF-treated samples versus irradiated samples (**Supplementary Fig. 9b**). Collectively, this study demonstrates Squidiff’s capability to predict cell-type-specific responses to drug treatments and reveals the effects of G-CSF on irradiated blood vessel systems.

## Discussion

In this study, we introduce Squidiff, a conditional diffusion model with a semantic encoder designed to predict transcriptomic changes across diverse cell types in response to a spectrum of environmental changes. Squidiff provides a robust framework for modeling cell development and responses on single-cell transcriptomic data, offering insights into cell differentiation and therapeutic responses, and positioning itself as a valuable tool in precision medicine. Its conditional design allows flexible control of experimental factors, while the integration of rFCFP embeddings enables prediction for previously unseen drugs.

We applied Squidiff to blood vessel organoids to investigate the effects of neutron irradiation and G-CSF treatment, a radioprotective intervention. As vascular integrity plays a central role in maintaining blood flow, nutrient distribution, and metabolic homeostasis, radiation-induced endothelial damage poses major risks in deep space missions^53–55^. A key advantage of Squidiff is its ability to infer dynamic, cell-type-specific responses from single timepoint datasets, thus circumventing the high costs and sample heterogeneity challenges with multi-timepoint single-cell sequencing. Despite sequencing being performed one week post-irradiation, Squidiff reconstructed early injury-phase transcriptomic changes, identifying temporal gene responses as potential radioprotective targets. Our analysis also highlights G-CSF’s therapeutic potential in vascular protection, providing insights into strategies to enhance its delivery efficacy and therapeutic index *in vivo*. These findings open avenues for future studies exploring other protective agents in space medicine and related fields.

Despite these promising results, Squidiff has several limitations. The training process involves introducing Gaussian noise into the data, resulting in prolonged training times. Furthermore, diffusion models generally require more computational resources compared to other generative frameworks, such as VAEs or generative adversarial networks. These challenges underscore the need for optimizing training protocols and more efficient implementations. Additionally, the current assumption of linearity in semantic variables may only provide approximate predictions in highly complex scenarios, requiring future refinement.

In the future, improving Squidiff’s scalability and computational efficiency will be essential to broadening applications to large-scale datasets and to establishing Squidiff as a more generalizable foundation model. While Squidiff has demonstrated strong performance in identifying cell-state transitions and perturbation responses, further validation using *in vivo* models would strengthen its translational potential. Moreover, extending Squidiff to integrate multi-modal omics data, including proteomics, epigenomics, and spatial information, may further enhance its predictive capabilities and enable the discovery of novel regulatory mechanisms governing cell fate decisions.

## Acknowledgments

We thank Stephen Quake and Konpat Preechakul for insightful discussions on virtual cells and the diffusion model, and Sabrina Wang, Jong Ha Lee, and Rayna Batya-Leia Berris for their assistance with single-cell RNA sequencing analysis and cell culture. We also appreciate the staff at Columbia’s Center for Radiological Research for their support in operating ionizing radiation equipment. This study utilized resources from the Herbert Irving Comprehensive Cancer Center Confocal, Specialized Microscopy Shared Resource, the Genomics and High Throughput Screening Shared Resource, partially funded by the NIH/NCI Cancer Center Support Grant P30CA013696. The Columbia IND Neutron Facility was developed under NIAID Grant U19 AI067773. We gratefully acknowledge funding support from the Translational Research Institute for Space Health (TRISH/NASA) (RAD0104 and NNX16A069A to G.V.-N. and K.W.L.).

## Author Contributions Statement

K.W.L., E.A. and J.Z. conceived the study and provided overall supervision of the study. S.H., Y.Z., D.N.T. designed the blood vessel organoids study and performed the experiments. S.H. and H.Y. designed and developed model. S.H., Y.Z., D.N.T., H.Y., Z.Z., C.X., S.C. analyzed and interpreted data. G. G. performed irradiation experiments. R.T. and G.V.N. provided additional supervision. S.H., Y.Z., D.N.T., H.Y., J.Z., E.A. and K.W.L. wrote the paper. All authors reviewed, contributed to and approved the paper.

## Methods

### Ethics Statement

All procedures involving human induced pluripotent stem cells (iPSCs) were conducted in accordance with institutional and federal ethical regulations, with guidance from the Columbia Stem Cell Initiative (CSCI). The established WTC11 hiPSC line was obtained from B. Conklin at the Gladstone Institutes under a material transfer agreement (to G.V.-N.).

Diffusion probabilistic models (DPMs) are a class of latent variable generative models that iteratively transform data into noise and then reverse this process to reconstruct the original data. A DPM consists of the forward process, the reverse process, and the sampling procedure. The model learns to reverse the diffusion processes step-by-step, capturing complex data distributions. A variant of DPM, the denoising diffusion implicit model (DDIM), further modifies the reverse process to be deterministic, improving sampling efficiency by eliminating the need for iterative sampling steps. DPMs and DDIMs have proven to be powerful methods for generating new data in various domains, such as images^1^, videos^2^, and protein structures^3^ in biomedicine. These models have also shown high potential in single-cell reconstruction and inference^4^, although their actual implementation in genetics is still limited. This limitation may be due to the unique challenges of applying DPMs to generate gene expression data, particularly the need for semantic manipulation and meaningful interpretation of the generated expression profiles. Recent advances in combining DPMs and variational autoencoders (VAEs) have demonstrated effective representation learning in the image domain^5^. Building on these advancements, we adapted the diffusion autoencoder model^5^, also a conditional denoising diffusion implicit model (DDIM) model for single-cell gene expression prediction under various perturbations. The goal is to learn a semantically rich latent space that allows smooth interpolation while maintaining the reconstruction capability that the diffusion model excels at.

### Squidiff model

Similar to other diffusion autoencoder models, the goal of Squidiff is to model the target distribution of perturbed gene expression by learning a denoising process at varying noise levels. The diffusion process, reverse process, and the semantic encoder are the three major components of the Squidiff model (**Fig. 1B**). Specifically, biologically meaningful information, such as cell type, environmental changes, and disease states, are encoded via the semantic latent variable *z*_sem_, as these perturbations are assumed to be inherently reflected in the transcriptomic data under given conditions. On the other hand, stochasticity is controlled via the random variable *x*_*T*_.

### Diffusion process

Given input data, the gene expression vector in an individual cell is denoted as *x*_0_. We first define the Gaussian diffusion process, which increasingly adds noise to *x*_0_ at time *t, t* ∈ {0,1,2,3, …, *T*}. The forward diffusion process is defined as:

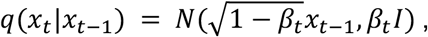

where *β*_*t*_ are hyperparameters representing the noise levels. This leads to 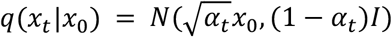, and α_*t*_ = Π^*t*^_*s*=1_ (1 − *β*_*s*_).

In our implementation, these parameters are defined as follows:

- *β*_*s*_ values are generated linearly spaced between 0.001 and 0.01
- The cumulative product of *β*_*s*_ values is computed to obtain *α*_*t*_
- The square root of these cumulative products and their complements are used to scale *x*_0_ and the noise, respectively.

The forward diffusion step, which adds noise to the gene expression vector, follows the equation: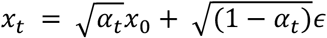, where *ϵ* is Gaussian noise.

#### Reverse process

The reverse process of DPMs aims to learn the noise distribution *p*(*x*_*t*−1_|*x*_*t*_), which is an intractable and complex distribution. Fortunately, we know from the diffusion process that *p*(*x*_*t*−1_|*x*_*t*_, *x*_0_) is a Gaussian distribution. In this regard, the diffusion model firstly estimates the clean data “*x*_0_” using the mean of distribution *p*(*x*_0_|*x*_*t*_), denoted as *μ_θ_*(*x*_*t*_, *t*):

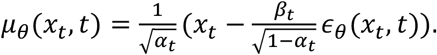

Here *ϵ_θ_*(*x*_*t*_, *t*) is the noise predicted by the denoiser, i.e., a neural network parameterized by *θ*. Based on the estimation, the diffusion model will sample *x*_*t*−1_ from *p*(*x*_*t*−1_|*x*_*t*_ = *x*_*t*_, *x*_0_ = *μ_θ_*(*x*_*t*_, *t*)).

The most famous denoising diffusion implicit model (DDIM) suggests the following *deterministic* generative process:

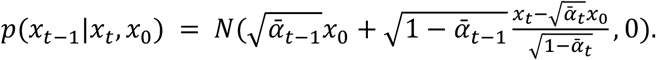

This ensures a deterministic transformation from *x*_*t*_ back to *x*_*t*−1_. In order to guide the denoising process via the semantic latent variable *z*_*sem*_, the denoising neural network in Squidiff is conditioned on the variable, i.e. *ϵ*_*θ*_(*x*_*t*_, *t, z*_*sem*_), and it predicts *x*_0_ by

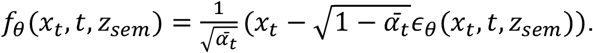

This ensures that the predicted *x*_0_ is conditioned on the environments represented by *z*_*sem*_, which is derived from a semantic encoder (see below) capable of capturing complex environmental features while performing dimension reduction (**Supplementary Fig. 1A**).

In summary, the conditional DDIM decoder takes *z* = (*z*_*sem*_, *x*_*T*_) as input, with the entire reverse process described as:

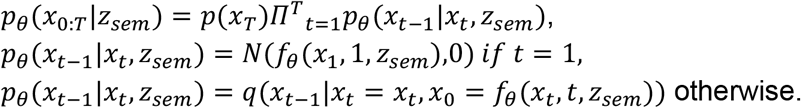

#### Semantic encoder

The goal of the semantic encoder is to provide *z*_*sem*_ for the reverse process to generate transcriptomics conditioned on *z*_*sem*_. The semantic encoder is designed to capture high-level semantic features from the input data, which are then used to condition the reverse diffusion process, ensuring that the generated transcriptomics data adheres to the desired semantic characteristics.

Our semantic encoder is implemented as a multi-layer perceptron (MLP)-based encoder designed for processing structured input data with optional auxiliary features like drug-related information (see sections below). The architecture consists of sequential linear transformations with batch normalization and ReLU activation, ensuring efficient feature extraction and transformation (**Supplementary Fig. 1A**). The semantic encoder is trained as part of the overall model. During training, the semantic encoder takes the input data and produces *z*_*sem*_, which is then used in the reverse diffusion process to conditionally generate the transcriptomics data. The training process ensures that the generated data is semantically coherent and adheres to the desired characteristics specified by *z*_*sem*_, which is equal to Enc(*x*_0_).

#### Sinusoidal position embeddings

To incorporate temporal information into the model, we use sinusoidal position encoding Ψ(*t*) for time step *t*, which allows the model to represent time as a continuous function rather than a discrete step. The core of this embedding lies in its frequency scaling. It applies an exponentially decaying function that controls how the frequencies are distributed, ensuring that both short-term and long-term dependencies are encoded effectively.

#### Training

The training process optimizes the objective function: 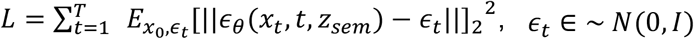, where the noise predictive function *ϵ*_*θ*_(*x*_*t*_, *t, z*_*sem*_) is modeled using a multi-layer perceptron (MLP) architecture conditioned on both the timestep *t* and semantic features *z*_*sem*_. The model architecture consists of an initial linear transformation followed by a sequence of MLP blocks incorporating timestep embeddings and latent semantic representations. Each MLP block performs a linear transformation followed by SiLU activations. The architecture incorporates a conditional residual mechanism where timestep embeddings and semantic features from an auxiliary encoder are added to the transformed representation, enhancing the model’s ability to capture temporal dependencies and contextual information (**Supplementary Fig. 1A**). The MLP is trained using the Adam optimizer with a learning rate of 1 × *e*^−4^.

#### Encoding drug structure information

Squidiff is primarily trained on transcriptomic data. However, in cases where transcriptomic data alone may not be sufficient — such as predicting responses to unseen drug perturbations — we introduce an adapter module inspired by PRNet^6^ (**Fig. 3H**). This module is not involved in the training process but is used to encode SMILES-based drug structure and dosage information into a latent representation. This drug-encoded representation is then concatenated with the original semantic latent variable *z*_*sem*_, forming an extended representation *z*_*sem*_′. In future work, Squidiff could incorporate additional environmental factors as needed to enhance its predictive capability.

### Simulation of single-cell RNA sequencing data

We simulated the single-cell RNA sequencing data using Splatter and its simulation model Splat. Splatter is an R Bioconductor package that provides a unified interface for multiple published simulation methods, including its own Splat model, which generates synthetic scRNA-seq data that closely resembles real datasets^7^. Splat is a gamma-Poisson hierarchical model, where the mean expression level for each gene is sampled from a gamma distribution, and the observed count for each cell is drawn from a Poisson distribution. Splat accounts for expression outliers and imposes constraints on variance to better mimic real scRNA-seq data characteristics. To obtain simulated scRNA-seq data from three distinct groups, we used Splatter and Splat with their default parameter settings.

### Data preprocessing and quality control

To ensure robust downstream analysis, single-cell RNA sequencing data underwent quality control:

- Cells with fewer than 1,000 expression counts or more than 20% mitochondrial gene expression were excluded.
- Genes expressed in fewer than three cells were filtered out.
- Potential multiplets were removed by excluding cells with over 10,000 detected genes.
- Mitochondrial and ribosomal genes, often associated with stress responses, were excluded.

Following quality control, the gene count data were normalized and log-transformed to correct for sequencing depth variability. Analyses then focused on highly variable genes and specific genes of interest, using Scanpy V1.10.1^8^ for processing.

### Squidiff training tasks

1. *Prediction of iPSC differentiation:* To model induced pluripotent stem cell (iPSC) differentiation, we selected the top 203 most variable genes, balancing biological significance with computational efficiency. Data collected on days 0 and 3 were used as the training datasets, while data on days 1 and 2 served as the testing datasets, respectively, resulting in 2,400 cells for training and 2,400 cells for testing. Gaussian noise was added to training data, and PCA plots confirmed 1,000 diffusion steps as optimal (**Supplementary Fig. 2A**). After training, the semantic variable for differentiation was computed by the mean difference between the latent representations at day 3 and day 0”:*Δz* = *E*[*z*^3^_*sem*_ ] − *E*[*z*^0^ _*sem*_]. The model then generated interpolated scRNA-seq representations using 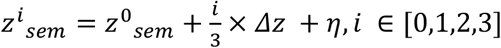, where η is a Gaussian noise term. This linear interpolation allowed the generation of synthetic single-cell RNA-seq profiles that smoothly transitioned from day 0 to day 3.
2. *Prediction of two non-additive genes perturbations*: The processed data were grouped into four categories: control, *PTPN1*+control, *ZBTB25*+control, and *PTPN12*+*ZBTB25*. The dataset was split such that the first three groups were used for training, while the last group served as the testing dataset. The diffusion process was set to 1000 steps. After training, the semantic variables for the trained groups were inferred, and the gene perturbation-specific variables were computed as follows:
  a. *Δz*^*PTPN*12^ = *E*[*z*^*PTP*12+*c*o*ntr*o*l*^_*sem*_] − *E*[*z*^*c*o*ntr*o*l*^_*sem*_]
  b. *Δz*^*ZBTB*25^ = *E*[*z*^*ZBTB*25+*c*o*ntr*o*l*^_*sem*_] − *E*[*z*^*c*o*ntr*o*l*^_*sem*_] To generate the predicted two-gene perturbed single-cell RNA sequencing data, we manipulated the latent representation by applying the learned perturbation variables: *z*^*PTPN*12+*ZBTB*25^ = *z*^*c*o*ntr*o*l*^_*sem*_ + *Δz*^*PTPN*12^ + *Δz*^*ZBTB*25^. These modified semantic representations were then used as conditions to simulate the transcriptomic changes induced by the combined perturbation of *PTPN12* and *ZBTB25*.
3. *Prediction of drug perturbation:* We selected highly variable genes along with some specific genes of interest, such as oligodendrocyte markers. The training data included oligodendrocytes, tumor cells, and myeloid cells, with conditions where these cells were either untreated (vehicle) or treated with etoposide. Additionally, myeloid cells treated with panobinostat, R04929097, tazemetostat, ispenisib, and Ana-12 were included in the training dataset. This setup ensured that two of the cell types (Oligodendrocytes and tumor cells) had never been exposed to drug treatments other than etoposide during training. As a result, Squidiff was challenged to generalize its predictions to novel drug-cell interactions. Squidiff learned the latent representation of each drug by computing the drug-specific perturbation effect in the latent space:

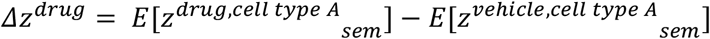

This formulation allows Squidiff to capture the distinct transcriptomic shifts induced by each drug, enabling it to predict the response of unseen cell types to novel drug treatments.
4. *Prediction of unseen drug perturbation:* While Squidiff demonstrates the capability to identify drug perturbations in unseen cell types, its predictive accuracy depends on the drug being trained on at least one cell type. When a drug is completely unseen during training, the model’s ability to predict its effect becomes limited due to the lack of prior exposure. To overcome this limitation, Squidiff incorporates an adaptor module that transforms drug-related molecular information, including the SMILES structure and dosage information, into an updated latent representation: *z*_*sem*_′ = *Enc*(*x*_0_, *rFCFP*), where *x*_0_ represents the baseline single-cell expression profile, and *rFCFP* denotes the fingerprint representation of the drug compound. By integrating these chemical and dosage features, Squidiff can generalize its predictions to completely unseen drugs. To evaluate this approach, we tested Squidiff using the sci-Plex3 dataset, performing two types of data splits: a. Random Split, where the dataset is randomly divided into training and testing sets. b. Unseen Drug Split, where drugs present in the test set were entirely absent from the training set, simulating a real-world scenario where the model must predict responses to completely novel compounds.

Squidiff was developed and tested with one NVIDIA H100 80GB HBM3. The time to process a dataset with 8000 cells and 500 genes takes about 15 minutes. While a high-performance GPU is recommended for efficiency, Squidiff can also run on lower-end GPUs, though with longer runtimes.

### Pseudotime analysis for single-cell RNA data

To further validate whether the predicted scRNA-seq data from day 0 to day 3 with Squidiff is biologically meaningful, we performed pseudotime analysis on the predicted dataset. The pseudotime values, as shown in **Fig. 2E** and **Supplementary Fig. 2D**, were computed using Scorpius^9^, an unsupervised approach for inferring linear developmental trajectories from single-cell RNA sequencing data. These values assign an ordering to each predicted cell, allowing us to assess the progression of differentiation over time. The distribution of pseudotime values for each real group, as well as the identification of gene expression trends along the trajectory, aligned well with expected biological patterns, supporting the validity of Squidiff’s predictions. To further benchmark Squidiff against established trajectory inference methods: PAGA, Monocle, and Scorpius, along with a random ordering baseline. Additionally, we calculated the Jaccard index of the top differentially expressed genes from day 0 to day 3 across the pseudotime series identified by PAGA, Monocle, Scorpius, random ordering, and the actual time points used in Squidiff. This comprehensive evaluation demonstrated that Squidiff effectively models differentiation trajectories from scRNA-seq data with strong agreement to experimental benchmarks.

### Reprogramming and culture of human pluripotent stem cells (iPSCs)

We cultivate the wild-type iPSC line (WTC) using mTeSR plus medium (Stem Cell Technologies) on growth factor-reduced Matrigel-coated 6-well plates. The cells are usually passaged every 3-4 days based on the proliferation rate.

### Generation of blood vessel organoids

To model the vascular malformations seen in PS, we generated blood vessel organoids using a previously established protocol^10,11^. The embryoid body (EB) was first formed in Aggrewell^®^ from iPSCs obtained one day prior to the induction of differentiation (Day -1). On day 0, organoids were distributed to 96-well plates. CHIR and BMP4 were added to induce the differentiation of embryoids into the mesoderm germ layer. On day 3, vascular differentiation was induced in the organoids by applying VEGFA and forskolin. The differentiated organoids were then embedded in Matrigel^®^ on day 5 and cultured in a medium containing VEGFA, FGF2, and FBS. The organoids were fixed on day 11 and stained for light sheet microscopic imaging^12^. We stained for endothelial and mural cell markers (CD31^+^, PDGFR*β*+, *α*SMA+, respectively) to confirm the generation of a vascular network constructed by endothelial tubes and mural cells, with visualization of the spatial arrangement of these cells in the vasculature.

### Radiation exposure

On Day 5 of BVO generation, we performed radiation exposure with either 1 Gy neutrons or 4 Gy photons, and the BVOs were cultured until Day 11. Neutron irradiations were performed at the Columbia University Radiological Research Accelerator Facility (RARAF), using an accelerator-based neutron irradiator^13,14^, as described previously^15^. The system was designed to mimic the neutron energy spectrum from an Improvised Nuclear Device^13^ but has also been used to model the high linear energy transfer components of galactic cosmic radiation^16^. Dosimetry was performed on the day of irradiation, as previously described, using a custom-built Tissue Equivalent Proportional Counter (TEPC), which measured the total dose, and a compensated Geiger–Mueller dosimeter, which had a very low response to neutrons and thus measured only the photon component. Photon irradiations were performed at Columbia University’s Center for Radiological Research using a Gammacell 40 137Cs irradiator (Atomic Energy of Canada Ltd) with a dose rate of 0.68 Gy/min for varying low-energy doses (i.e., 4 Gy).

### Drug treatment

Following radiation exposure, the medium from the blood vessel organoids was collected and replaced with a fresh vessel organoid medium. G-CSF at a concentration of 80 ng/mL was then administered to the samples immediately.

### Cytokine testing

Culture medium was collected from irradiated blood vessel organoids (BVOs) on days 0, 3, and 6 post-radiation to assess cytokine levels. TNF-*α* and IL-1*β* concentrations were quantified using uncoated ELISA kits from Thermo Fisher Scientific (Human TNF-*α* Uncoated ELISA Kit, Cat# 88-7346-22; Human IL-1*β* Uncoated ELISA Kit, Cat# 88-7261-88). For both assays, 96-well microplates were coated overnight at 4 °C with 100 µL per well of capture antibody diluted in coating buffer according to the manufacturer’s protocol. Plates were then washed three times with wash buffer (prepared by diluting the provided concentrate with distilled water) and blocked with 200 µL per well of ELISPOT Diluent for 1 hour at room temperature. Standard curves were generated by serial dilution of the cytokine standards: TNF-*α*, ranging from 500 to 7.8 pg/mL, and IL-1*β*, from 250 to 3.9 pg/mL. Irradiated BVO culture medium samples (100 µL per well) and standards were added in duplicate to the wells and incubated for 2 hours at room temperature with gentle shaking. After five washes, 100 µL of detection antibody was added and incubated for 1 hour, followed by 100 µL of Avidin-HRP for 30 minutes, with additional washing steps between each reagent. Colorimetric detection was performed by adding 100 µL of TMB substrate solution to each well and incubating for 15 minutes in the dark at room temperature. The reaction was stopped by adding 50 µL of 2 N H_2_SO_4_, and absorbance was measured at 450 nm with a reference wavelength of 570 nm using a microplate reader.

### Whole-mount Immunocytochemistry

The blood vessel organoids were fixed with 4% paraformaldehyde overnight at 4°C, followed by four PBS washes (15 min each). Samples were permeabilized and blocked with blocking buffer (1.5 mL FBS, 0.5 g BSA, 250 µL Triton X-100, 250 µL Tween 20, and 47.5 mL PBS in 50 mL total volume) for 2 hours at room temperature.

Primary antibodies were diluted in blocking buffer as follows:

Rabbit anti-human CD31 (1:400, Cat# ab28364, Abcam);

Goat anti-human *α*-smooth muscle actin (1:400, Clone EPR5368, Cat# ab320058, Abcam); Samples were incubated with primary antibodies overnight at 4°C.

Following six PBS-T washes (1 hour each), secondary antibodies were applied:

Alexa Fluor 488 Donkey Anti-Rabbit IgG F(ab’)2 (1:1000, Cat# 711-546-152, Jackson ImmunoResearch);

Alexa Fluor 647 Donkey Anti-Goat IgG F(ab’)2 (1:1000, Cat# 705-606-147, Jackson ImmunoResearch);

Secondary antibodies were incubated overnight at 4°C in the dark. Nuclei were stained with DAPI (1 µg/mL, Cat# D1306, ThermoFisher) during the final wash. After six PBS-T washes, samples were cleared overnight with tissue-clearing reagent (RI=1.49) before mounting.

### Confocal imaging

The blood vessel organoids were permeabilized with blocking buffer (1.5 mL FBS, 0.5 g of BSA, 250 μL of Triton X-100, 250 μL of Tween 20, and 47.5 mL of PBS in 50 mL buffer) and incubated with primary antibody overnight at 4°C. The samples were then washed with PBS-T solution and embedded in a mounting medium. Whole-mount 3D imaging was performed using a Nikon confocal microscope with a 10× objective. Tile scanning and Z-stack acquisition were used to capture the complete blood vessel organoid structure.

### Single-cell sequencing

Single-cell RNA sequencing was performed in BVOs on day 11. Blood vessel organoids (72 organoids for each condition) were dissociated into single cells using enzymatic digestion with a dissociation buffer containing trypsin or collagenase. The dissociated cells were counted using an automated cell counter, and cell viability was assessed using trypan blue exclusion. Single-cell suspensions were then loaded into a microfluidic device, such as the 10x Genomics Chromium Controller, to capture individual cells in nanoliter-scale droplets containing reagents for cell lysis and reverse transcription. Unique molecular identifiers (UMIs) and cell barcodes were incorporated to enable accurate identification and quantification of transcripts from individual cells. After reverse transcription, the cDNA was amplified using polymerase chain reaction (PCR) and purified. Sequencing libraries were constructed from the amplified cDNA using a standard library preparation kit and indexed with unique sample barcodes for multiplexing. The libraries were sequenced on a high-throughput platform, Illumina NovaSeq, to generate paired-end reads optimized for comprehensive coverage and accurate quantification of gene expression.

The raw sequencing data were processed using the Cell Ranger pipeline (version 6.1.2) to demultiplex reads, collapse duplicates based on UMIs, align reads to the GRCh38 reference transcriptome, and quantify gene expression levels for each cell. Quality control steps removed low-quality cells and potential doublets, and the high-quality single-cell RNA-seq data were analyzed for normalization, clustering, differential expression, and trajectory inference to elucidate the transcriptomic changes in response to different treatments.

## Data Availability

The public dataset of single-cell RNA-sequencing of iPS cells differentiating towards endoderm^17^ was downloaded from https://zenodo.org/records/3625024#.Xil-0y2cZ0s. The dataset for non-additive gene perturbation^18^ was downloaded from GSE133344. The dataset for drug treatment on melanoma cells^19^ was downloaded from https://www.research-collection.ethz.ch/handle/20.500.11850/609681. The public dataset of drug screening on glioblastoma^20^ was downloaded from accession GSE148842. The sci-plex3 dataset^21^ was downloaded from GSM4150378. The raw single-cell sequencing data of BVO generated is available in figshare: https://doi.org/10.6084/m9.figshare.27948633.

## Code Availability

The Squidiff package and code to reproduce the results in this manuscript are available on the GitHub repositories:

https://github.com/siyuh/Squidiff and https://github.com/siyuh/Squidiff_reproducibility. The code is also deposited at Zenodo (https://doi.org/10.5281/zenodo.15061773)^22^.

